# Managing human mediated range shifts: understanding spatial, temporal and genetic variation in marine non-native species

**DOI:** 10.1101/2021.07.06.451303

**Authors:** Luke E. Holman, Shirley Parker-Nance, Mark de Bruyn, Simon Creer, Gary Carvalho, Marc Rius

**Affiliations:** School of Ocean and Earth Science, National Oceanography Centre Southampton, University of Southampton, United Kingdom; Nelson Mandela University, Gqeberha (Port Elizabeth), South Africa; South African Environmental Network (SAEON) Elwandle Coastal Node, Gqeberha (Port Elizabeth), South Africa; The University of Sydney, School of Life and Environmental Sciences, Australia; Molecular Ecology and Evolution Group, School of Natural Sciences, Bangor University, United Kingdom; Department of Zoology, Centre for Ecological Genomics and Wildlife Conservation, University of Johannesburg, South Africa

**Keywords:** ascidians, biodiversity, environmental DNA, non-native species, range shifts

## Abstract

The use of molecular methods to manage natural resources is increasingly common. However, DNA-based methods are seldom used to understand the spatial and temporal dynamics of species’ range shifts. This is important when managing range-shifting species such as non-native species (NNS), which can have negative impacts on biotic communities. Here we investigated the range-shifting NNS *Ciona robusta, Clavelina lepadiformis, Microcosmus squamiger* and *Styela plicata* using a combined methodological approach. We first conducted non-molecular biodiversity surveys for these NSS along the South African coastline, and compared the results with historical surveys. We detected no consistent change in range size across species, with some displaying range stability and others showing range shifts. We then sequenced a section of cytochrome c oxidase subunit I (COI) from tissue samples and found genetic differences along the coastline but no change over recent times. Finally, we found that environmental DNA metabarcoding data showed broad congruence with both the non-molecular biodiversity and the COI datasets, but failed to capture complete incidence of all NSS. Overall, we demonstrated how a combined methodological approach can effectively detect spatial and temporal variation in genetic composition and range size, which is key for managing biodiversity changes of both threatened and NSS.

## 1. Introduction

Biodiversity is undergoing a global redistribution as a result of human influence, with species increasingly found in environments outside their previously reported geographic range [1]. Contemporary climate change is causing species to shift their ranges to accommodate novel environmental conditions [2, 3], and human-mediated species introductions dramatically increase the range of non-native species (NNS) [4-6]. This exposes species to abiotic conditions and biotic interactions that are different to those experienced in native habitats. Such changes in distribution can result in a dramatic increase or decrease in population size, or may have a limited detectable immediate effect [1, 7]. Understanding these responses is important to answer fundamental ecological and evolutionary questions about changing biotic communities, but also for natural resource managers when predicting changes in ecosystem services and natural capital [1, 8].

Global biodiversity loss has consistently been shown to reduce ecosystem function, and in turn affects the provision of ecosystem services [9, 10]. A key driver of biodiversity loss is the introduction of NNS [11], which also imposes a substantial global economic cost [12] and has a dramatic impact on public health [13, 14]. In the marine environment the majority of NNS introductions are associated with transoceanic shipping [5, 15, 16] and therefore, major ports and harbours are hotspots for NNS. Once a species is introduced to these sites, secondary spread can be facilitated by smaller recreational vessels, marinas and marine infrastructure surrounding these major harbours [17, 18]. Considering the increasing number of yearly NNS introductions [6, 19], improving our understanding of how range shifts of NNS occur through time and space is critical in the design of effective management and mitigation responses.

Natural resource managers have finite budgets and limited information when making decisions simultaneously on a number of NNS with variable or unknown impact [20, 21]. For each NNS, managers can attempt to eradicate a population, make efforts to avoid any further expansion into new areas, or acknowledge that control is not possible and work on mitigation strategies [22, 23]. These limited options are compounded by the vast costs associated with control or eradication, and even when control methods may be possible, they might be politically or publicly unacceptable [24, 25]. Furthermore, control measures can be unsuccessful because of incomplete eradication of the target species or ongoing species reintroductions [20, 26, 27]. Consequently, managers frequently take no action to control NNS or act only when evidence for both presence and substantial impact has been gathered [28]. It is therefore beneficial to develop tools that provide researchers and managers with information to facilitate decision-making. Genetic tools can complement existing methods for assessing NNS range shifts by providing information that would be unfeasible or impossible to produce otherwise [29].

Even when NNS can be unambiguously identified, it can be difficult to determine when and where they were first introduced into a region, (for example see Hudson *et al*. [30]). Since eradication or control efforts are improved by early detection [31], methods with high sensitivity are needed to increase the likelihood of successful management outcomes. One such method is the isolation of DNA from environmental samples (environmental DNA or eDNA) such as water or sediment for the detection of organisms. Studies have demonstrated that the amplification of DNA barcode regions from eDNA (i.e. eDNA metabarcoding) can be used to detect marine NNS [32-35] and that it is a sensitive and accurate method for biomonitoring [36, 37]. However, eDNA surveys are rarely used in conjunction with existing methods to detect NNS range shifts, and eDNA metabarcoding can validate, endorse, or highlight flaws, in current biodiversity management strategies.

During a range expansion, understanding if there was a single NNS introduction event or multiple simultaneous introductions is valuable for managers to target possible source regions, and to effectively manage introduction vectors (e.g. ballast waters). As NNS spread across the new region, understanding if expansions are due to local spread or introductions from distant regions is useful to target containment efforts. Finally, after eradication efforts have been conducted, understanding if the reappearance of NNS is due to incomplete eradication or a secondary reintroduction is of value for effective management into the future. The sequencing of DNA isolated from NNS has previously identified the source of NNS [38, 39], provided evidence of multiple introductions [40] and tested if post eradication invasions are a result of incomplete eradiation or reinvasion [41]. Cumulatively, these studies have demonstrated the value of DNA evidence for the management of NNS. Furthermore, observations from both laboratory [42, 43] and field studies [44-49] have shown that eDNA can provide population genetics inference, but very little work has used this approach to study NNS [50].

Here we combined eDNA metabarcoding, mitochondrial gene sequencing, and non-molecular biodiversity surveys to study four NNS that are directly relevant to marine natural resource managers. First, we evaluated if the NNS shifted their ranges over decadal time scales and compared each range shift to historical data. Secondly, we evaluated changes in genetic diversity and haplotype composition for each NNS between two sampling occasions across the sampled coastline. Finally, we examined how spatial genetic variation data can inform the management of range shifting species by comparing eDNA metabarcoding data to biodiversity survey and mitochondrial DNA sequence datasets.

## 2. Methods

### (a) Fieldwork and historical biodiversity data

The coastline of South Africa is an ideal system to study range shifting species and their management. South Africa has been subject to intense human impact and many species invasions have been documented across the three environmentally varied coastal ecoregions [51-53]. Moreover, rapid assessment survey (a non-molecular biodiversity survey technique) [54] data has been previously collected and mitochondrial sequence data have been generated for NNS along the entire coastline [52]. Furthermore, historical data are available for a range of relevant species [55-57] providing an insightful opportunity to conduct a spatial and temporal analysis of range expansions. Here, we selected twelve human impacted sites and conducted surveys (see details below) between October and November 2017. The sampled sites were the 11 sites previously sampled in 2007 and 2009 by Rius *et al*. [52], which included all major harbours and a number of marinas, and a new marina constructed post 2009 (Figure 1a with full details in Supplementary Table 1). Collectively, the sites encompass the main introduction points for marine NNS into the South African coastline.

**Figure 1.**
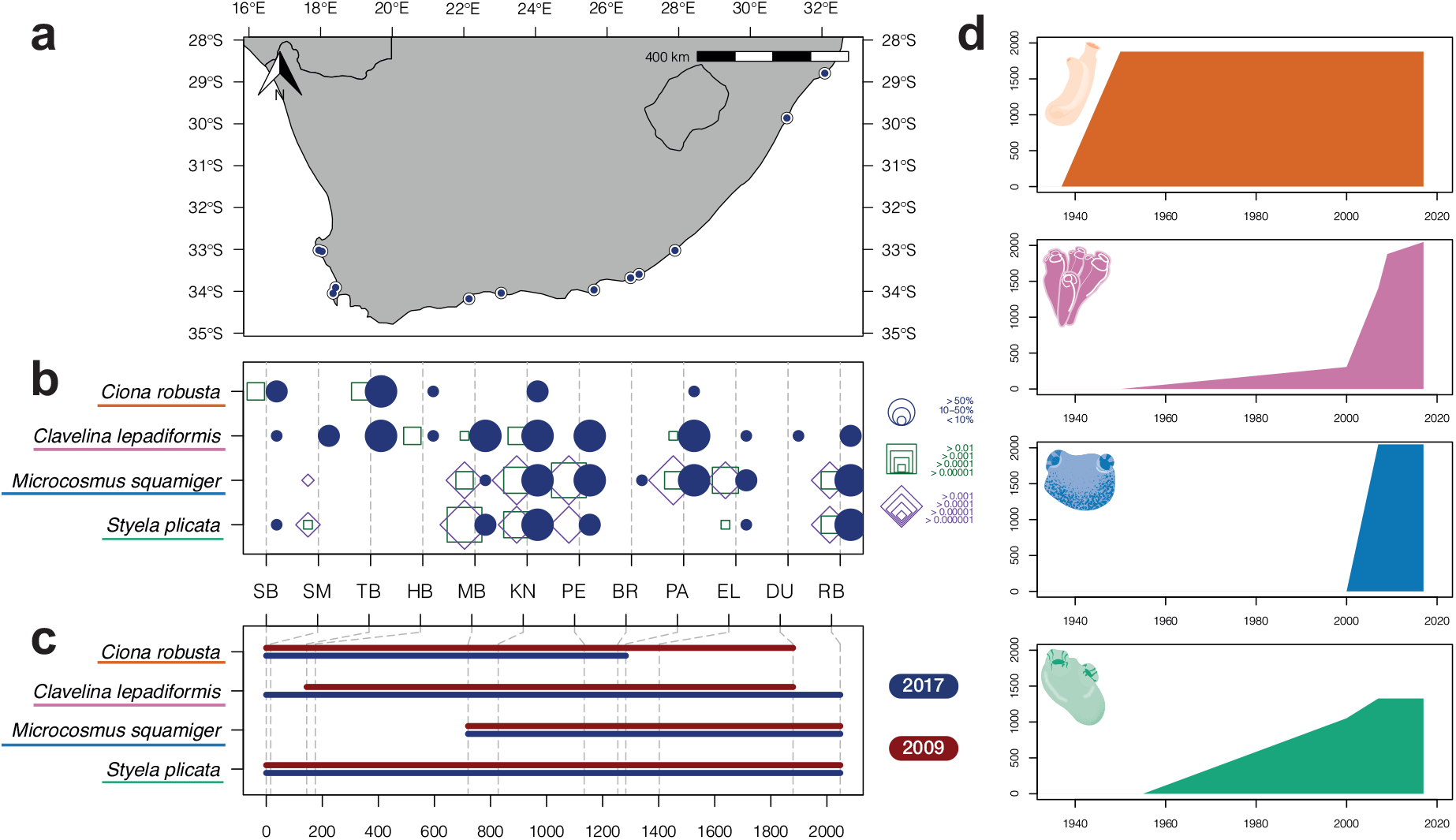
**a** Map depicting the coastline of South Africa, sampling sites are shown as blue points, full details in Supplementary Table 1. **b** Bubble plot showing incidence of four non-native ascidians across the sampling sites shown in the map from west to east. Blue bubbles show results of rapid assessment surveys and square outlines show the results of eDNA metabarcoding surveys conducted concurrently. Results from COI are shown with green squares and 18S shown with purple squares, the size of each point or square shows the comparative density. Site codes correspond with sites as detailed in Supplementary Table 1. **c** Line plot showing range extent over the surveyed coast for 2009 (dark red) rapid assessment surveys from Rius *et al*. [52] and surveys conducted in 2017 presented here (blue). The location of each site across the coastline is shown with grey dashed lines. **d** Historical maximum range extent for each of the featured species across the coastline of South Africa, y axis is kilometres of extent, x axis is year, colour indicates each of the species indicated according to labels in b and c.

At each sampling site a rapid assessment survey was conducted following Rius *et al*. [52], targeting non-indigenous ascidian species (Class: Ascidiacea). Ascidians are unique species for studying range expansions as they are successful invaders [58] and have a relatively short pelagic larval phase, meaning that long-distance dispersal can only be achieved through anthropogenic transport of species [59]. For each site, species abundance was ranked as absent (0%), scarce (< 10%), common (10-50%), or dominant (> 50%) based on observations of substrate coverage as in [52].

Rapid assessment survey data from 2007 and 2009 was sourced from Rius *et al*. [52] for the species of interest. Additionally, historical incidence data was extracted from several taxonomic publications [55-57, 60, 61]. These investigations are not an exhaustive survey of the coastline, but they provide valuable historical species incidence data over the last century and are therefore of value in gaining a broad understanding of range shifts over time.

### (b) Sample collection, DNA extraction and Sanger sequencing

Tissue samples were collected for species for which genetic data were available from the 2009 surveys (*Ciona intestinalis, Clavelina lepadiformis, Microcosmus squamiger* and *Styela plicata*). Samples were collected where sufficient numbers of individuals per species were present at a site to provide a reasonable estimate of genetic diversity (minimum 10 individuals), with 30+ individuals per species being the target at each sampling site. Organisms were sampled by hand, with no adjacent (within 0.3m) individuals collected, and dissected within six hours (see details of research permit in the Acknowledgements). For each sampled individual, approximately 10mm^2^ of tissue from around the syphons was dissected using tools decontaminated with 10% bleach solution (3.5% chlorine), except in the case of *C. lepadiformis* for which a single zooid was removed from the tunic and stored. Tissue samples were preserved in 100% ethanol and stored at ambient temperature during transportation, and then stored at -80°C in the laboratory until later DNA extraction. DNA from ascidian tissue samples was extracted using the Qiagen (Hilden, Germany) DNeasy Blood and Tissue Kit (96 Well Format) following manufacturer’s recommended protocol with one blank control per extraction run. The final DNA elution was performed using 200μl of Qiagen Buffer ATL. A section of the cytochrome c oxidase subunit I gene (COI) was sequenced for all tissue samples aiming to cover the entire section previously analysed in Rius *et al*. [52]. Each PCR contained 6μl of Applied Biosystems (Foster City, Califonia, USA) AmpliTaq GOLD 360 Mastermix, 1.8μl of oligonucleotide mix (5 μm concentration per primer), 1.2μl of undiluted template DNA and PCR quality water up to 12μl total reaction volume. The reaction conditions varied by primer set and are listed in Supplementary Table 2a. During preliminary trials a set of primers were designed and validated for *M. squamiger* (sequence details in Supplementary Table 2b), existing primer sets [62-64] were optimised for the remaining three species. Successful amplification was confirmed using gel electrophoresis and PCR products were cleaned using Applied Biosystems ExoSAP-IT Express following the manufacturer’s recommended protocol. Cleaned products were normalised to approximately 50ng/μl and 5μl of sample was added to each of 5μl of the forward or reverse primers (5μm) used in the initial PCR. These samples were sent for sequencing using the Macrogen Europe (Amsterdam, Netherlands) EZ-Seq service. Resultant chromatogram files were analysed using Geneious Prime (v2020.2.4) (Biomatters Ltd, Auckland, New Zealand). For each sequence the forward and reverse traces were aligned and sequences with ambiguities or failed reactions were re-sequenced from the initial PCR once and subsequently discarded if poor results persisted. The 764 COI sequences from Rius *et al*. [52] were added to the analysis and trimmed, truncated and aligned with the experimental data as follows. For each species, sequences were trimmed to remove primer binding and poor-quality regions and aligned using the Geneious Alignment Tool. Subsequently each alignment was manually checked to confirm complete alignment, and short sequences that did not overlap at all polymorphic regions or had ambiguous base calls were discarded.

### (c) Environmental DNA metabarcoding

Before each rapid assessment survey, surface seawater was sampled from the top 10cm for eDNA metabarcoding following Holman *et al*. [65]. Briefly, three replicate 400ml water samples were filtered on site with a 0.22 µm polyethersulfone enclosed filter. Filters were preserved with Longmire’s solution until DNA extraction following Spens *et al*. [66]. Data generated from these samples is presented in Holman *et al*. [65] with the aim of conservatively characterising whole community diversity. COI and ribosomal RNA (18S) data targeting metazoans [67, 68] was reanalysed as follows for accurate ascidian species detection. Primer regions were removed from forward and reverse reads using the default settings of Cutadapt (v2.3) [69]. Sequences were denoised and an ASV (amplicon sequence variant) by sample table generated using DADA2 (v1.12) [70] in R (v3.6.1) [71] with parameters as in Holman *et al*. [65]. Recent work has highlighted that different bioinformatic methods have an effect on the resolution of intra-specific variation of eDNA metabarcoding data [42, 45, 72]. Therefore in addition to the sequenced tissue samples and DADA2 methods outlined above, we reanalysed the COI data using the *unoise3* algorithm [73] as follows. Raw COI paired-end fastq data from Holman *et al*. [65] was merged using usearch (v11.0.667) [73] with the following parameters *-fastq_maxdiffs 15 -fastq_pctid 80*. Primer sequences were then stripped from each merged read using Cutadapt (v3.1) [69] under the default parameters and reads longer than 323 and shorter than 303 base pairs (±10 from the expected size of 313) were discarded. Reads from all samples were pooled, and singletons and reads with an expected error greater than 1 were discarded using vsearch (v2.15.1) [74]. The *unoise3* algorithm from usearch was then used to generate ASVs with *-unoise_alpha* set at 5 as recommended for resolving metazoan intraspecific variation with a COI fragment of 313 base pairs in length [45]. Sequences were then mapped back to the ASVs using the -u*search_global* function of vsearch with an *-id* parameter of 0.995 to produce an ASV by sample table.

To provide an initial taxonomic assignment all ASVs were compared using a BLAST (v2.6.0+) search with no limits on sequence similarity or match length to the NCBI *nt* database (downloaded 16^th^ May 2019). Taxonomic assignments were then parsed using a custom R function (*ParseTaxonomy*, DOI:10.5281/zenodo.4671710) with the default settings. The taxonomic assignments were subset to include only those with a hit to species in the class Ascidiacea. The following quality control steps were then applied to each dataset. The data was filtered to only retain ASVs that appeared in more than one replicate sample. For any ASVs detected in both the negative and experimental control samples, the maximum number of reads in the negative controls were subtracted from the experimental control samples. Reads were then divided by the total number of reads per sample and relative proportions were used in all subsequent analyses, technical replicates per site were averaged. The remaining ASVs were then taxonomically checked manually using the online National Centre for Biotechnology nucleotide BLAST search function against the *nt* databases (last accessed on 1^st^ October 2020) under default megablast parameters. For each ASV in the COI dataset, taxonomy was only assigned at species level if multiple, independent sequences had a match greater than 97% identity (with 100% coverage) with no other species within 97% of the target ASV. For the 18S dataset a 100% match (with 100% coverage) between the subject ASV and database sequences was required for taxonomic confirmation. Additionally, as some taxa within the same genera have near 100% similarity at the 18S region, taxonomy was only assigned to species if organisms from the same genera were in the database with at least 1 base pair between the query and species from the same genera. Following taxonomic annotation, ASVs assigned to the same species were merged for the distribution datasets. ASVs were kept separate for the haplotype reconstruction of the COI data.

### (d) Data manipulation and statistical analyses

Distances between sites along the coast were estimated by drawing a transect 1 km parallel to the coastline in Google Earth Pro (v7.3.2.5776) and calculating the distance between each pair of sites. The study area was plotted using the function *map* from the package *maps* (v3.3.0). Sequenced COI regions from 2009 and 2017 were aligned separately for each species using the Geneious aligner in Geneious Prime; alignments were truncated to include only overlapping regions. Sequences were manipulated using the *SeqinR* package in R (v4.2-5) [75]. Nucleotide and haplotypic diversity were calculated using the *nuc*.*div* and *hap*.*div* functions from the *pegas* package (v0.14) [76]. For each species an alignment was created between the tissue sampled COI sequences and the eDNA metabarcoding derived haplotypes. The region of overlap was extracted and used in subsequent analyses. Haplotype frequencies were calculated per site for the tissue derived sequences and the different bioinformatic analyses of eDNA metabarcoding data. Minimum spanning network haplotype maps [77] were created using the default settings of PopArt (v1.7) [78]. Analyses of molecular variance (AMOVA) were performed using the function *poppr*.*amova* from the *poppr* package (v2.8.6) [79]. AMOVA models were structured to analyse the effect of sampling year and sites for each species. All data analyses were conducted in R (v4.0.3) unless otherwise stated.

## 3. Results

### (a) Range shifts

Rapid assessment surveys found that non-native ascidians known to be broadly restricted to warmer waters (*M. squamiger* and *S. plicata*) [52] showed distributions principally limited to the southern and eastern coastlines (Fig.1b). In contrast *C. robusta* and *C. lepadiformis* were found along most of the coastline. We found no change across years in range extent for *M. squamiger* and *S. plicata*, a decrease in easternly range for *C. robusta* and an expansion of range both westerly and easterly for *C. lepadiformis* (Fig.1c). Historical total range extent data (Fig.1d) showed more recent increases in range for *C. lepadiformis* and *S. plicata* compared to *C. robusta* and *M. squamiger*. The COI and 18S eDNA metabarcoding data showed mixed results. There was good agreement between detections from eDNA and rapid assessment surveys in *M. squamiger* and *S. plicata* (see Fig.1b). However, 18S entirely failed to detect *C. robusta* or *C. lepadiformis*, and COI demonstrated a number of false-negative metabarcoding detections in these species (Fig.1b). For sites sharing detections from eDNA metabarcoding and rapid assessment surveys, eDNA metabarcoding data and field density estimates showed a non-significant relationship (18S p = 0.052, COI p = 0.297) (see Supplementary Note 1 for details).

### (b) Changes in genetic composition

A total of 1,320 sequencing reactions generated 660 bi-directionally sequenced COI sequences. After alignment and quality control, 541 samples remained with complete alignment and no missing site information, 88 for *C. robusta*, 261 for *C. lepadiformis*, 90 for *M. squamiger* and 102 for *S. plicata*. After combining the COI sequences with previously sequenced samples from 2009 [52], alignments were 626, 440, 635 and 599 base pairs in length for *C. robusta, C. lepadiformis, M. squamiger* and *S. plicata* respectively. Observed haplotype richness across both sampling years and all sites was highest in *M. squamiger* followed by *C. robusta, S. plicata and C. lepadiformis* (Fig. 2). There was no statistically significant difference between nucleotide or haplotype diversity between sampling years across all species (p > 0.05 in all cases, see Supplementary Note 2 for full model output and details). Additionally, AMOVA models found no significant differences between sampling years across all species (p > 0.05 in all cases, see Supplementary Table 3 for full model outputs), but significant differences between sampling sites within years (p < 0.05 in all species, see Supplementary Table 3 for full model outputs). In all species, the greatest proportion of the genetic variance was found between samples, then within sampling sites, followed by variance between sampling sites (Supplementary Table 3). As shown in Figure 2, haplotype frequencies agreed with the AMOVA analyses, showing stable patterns of genetic variation occurring between years and variation in haplotype frequencies across the study system (Figure 2).

**Figure 2.**
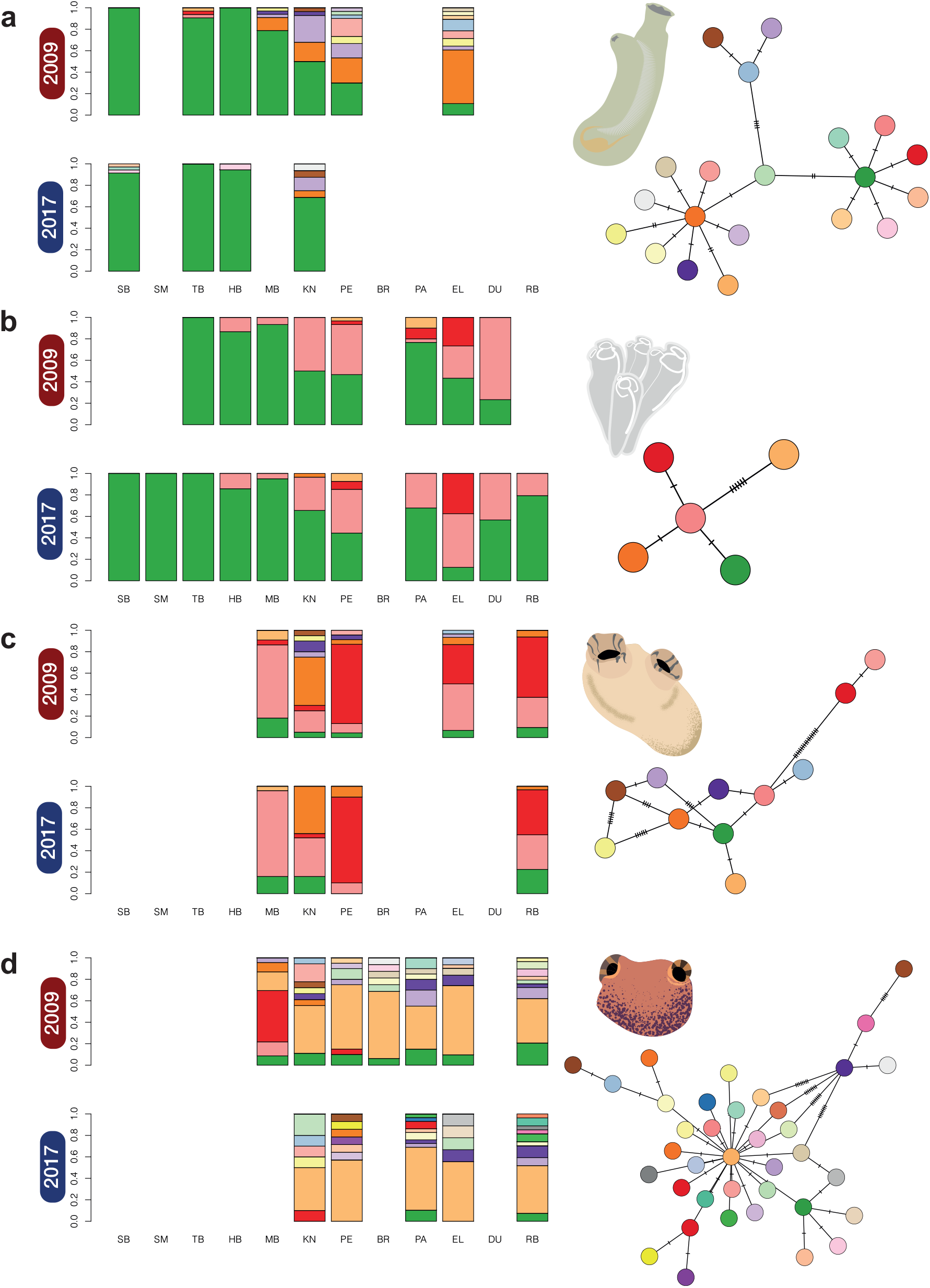
Mitochondrial DNA COI haplotype proportions for **a** *Ciona robusta* **b** *Clavelina lepadiformis* **c** *Styela plicata* **d** *Microcosmus squamiger* across the South African coastline. Results are shown for surveys conducted in 2009 and 2017 for each species, site abbreviations follow supplementary Table 1. Haplotype networks based on minimum spanning distance are shown for each species with colours matching the bar plot within species, the number of cross-hatches indicates the mutation steps between haplotypes.

After aligning the shorter sequences derived from eDNA metabarcoding data to the sequenced COI region, alignments were 191, 258, 289 and 286 base pairs in length for *C. robusta, C. lepadiformis, M. squamiger* and *S. plicata* respectively. Regardless of bioinformatic method and across species, the eDNA metabarcoding data did not recover all the haplotype sequences derived from tissue (Figure 3).

**Figure 3.**
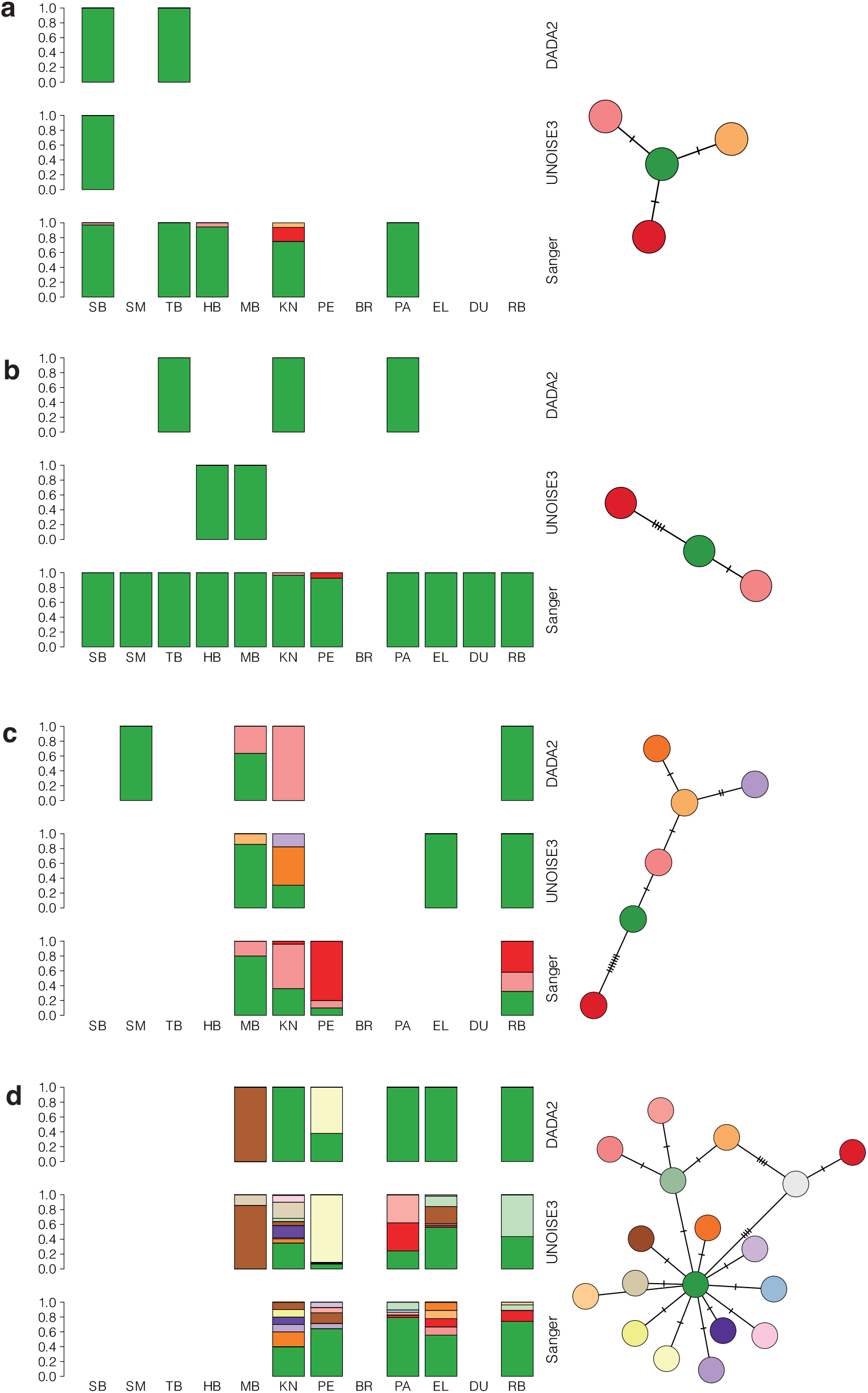
Haplotype proportions recovered using eDNA metabarcoding for **a** *Ciona robusta* **b** *Clavelina lepadiformis* **c** *Styela plicata* **d** *Microcosmus squamiger* across the South African coastline. Results are shown for analysis of COI eDNA metabarcoding data using the denoising software DADA2 and UNOISE3 for each species, site abbreviations follow supplementary Table 1. Haplotype networks based on minimum spanning distance are shown for each species with colours matching the bar plot within species, the number of cross-hatches indicates single nucleotide mutation steps between haplotypes.

## 4. Discussion

Here we found both losses and gains in range size across sampling years for four non-native ascidian species, with no consistent pattern emerging when introduction dates were compared. For all species we found haplotype variability across the study region but no significant change in genetic variation for almost a decade. Finally, eDNA metabarcoding data recovered broad NSS incidence trends and for some species was as accurate as non-molecular surveys. Most dominant haplotypes from tissue samples were detected with eDNA metabarcoding but fine scale genetic patterns could not be resolved using the eDNA metabarcoding data. Cumulatively, the evidence demonstrates that a suite of tools, including DNA and non-DNA biodiversity surveys can be used in combination to evaluate the role of genetic variation on range shifts and to inform natural resource managers.

Non-DNA biodiversity surveys found that *C. lepadiformis* expanded its range by 168.4 km since surveys in 2009, for an assumed rate of 21.1 km per year. This is in line with previous studies that found an average marine non-native spread rate of 44.3 km per year [80], with values of 16 km per year for tunicates, 30.0 km per year for barnacles and 20 km per year for a bryozoan species [81]. In contrast, we observed a range contraction for *C. robusta*, which was unexpected. There are limited studies showing range contraction in the introduced range for marine species. However, previous work has identified biotic resistance for invasions of several species in the genus *Ciona* [82, 83], and so it might be feasible for local species to have begun predating on *Ciona robusta* during the 80+ years it has been documented in South Africa (Fig.1d). A lack of any western increase in range for *M. squamiger* might be explained by the species inability to mature to reproductive age in the colder sea temperature on the western coast [52]. Further range expansions or contractions (eastwards for *M. squamiger* and east or westward for *S. plicata*) cannot be ruled out as observations of these species extended to the margins of the sampled area. It is important to note that the harbours and marinas in this study act as islands of suitable habitat, and the frequency of introductions outside these areas is relatively uncertain. Further surveys of surrounding hard benthic environments are required to understand the role of artificial environments across the coastal ecosystem. Overall, these patterns demonstrate that the spread of marine non-native species is not characterised by a continuous expansion of range, but rather by a complex picture of expansions and contractions in response to dynamic abiotic and biotic conditions.

A consistent pattern of genetic differentiation emerged across the studied species; significant differences across sampling sites and persistence of similar haplotypes across time (Fig. 2). Previous studies of temporal changes in genetic diversity of non-native ascidians have found some evidence for genetic differences over time [84, 85]. In contrast other work has found relatively stable genetic diversity over several years [86, 87]. In our study, the time between sampling occasions (i.e. 2009 and 2017) represents between four and 24 generations depending on the species [88-90]. Therefore, dramatic changes in haplotype frequencies could only be as a result of anthropogenic transfer of haplotypes between sites or changes in site frequencies in response to high mortality events (for example extreme weather events). These types of changes have been documented in ascidian species elsewhere [85, 91, 92], and a large number of NNS introductions have been documented in South African marinas and harbours supporting regional transfer of these organisms [52, 53]. It is therefore somewhat surprising that across four different species, all of which are known to be transported anthropogenically, there was little evidence of shifts in haplotype composition. Consequently, our results demonstrated that the studied NSS are well-established and are not subject to high levels of mortality or genetic bottleneck that may affect population viability. It may be that these well-established haplotypes impede newcomers to succeed and ultimately change the haplotype composition of the site.

We found that eDNA metabarcoding captured similar incidence data as rapid assessment surveys for some species, and performed poorly for others. Previous work has identified that NNS can be detected using eDNA metabarcoding [32, 34], but these surveys aimed at detecting any NNS rather than a specific set of target taxa. Several studies have identified that general target metabarcoding primers show lower reliability and sensitivity compared to species-specific quantitative PCR assays [93, 94]. Additionally, previous work has identified that in some cases different bioinformatic methods carry variable sensitivity [95], although this effect is fairly minimal in this dataset (see Supplementary Note 3). Managers should therefore be aware that general metabarcoding primers will perform well in the detection of some important NNS but others may be missed due to poor sensitivity. In cases when a list of priority species can be assembled, mixed DNA positive control samples or trials with aquaria of known composition (for example Holman *et al*. [96]) would provide information on which NNS might be overlooked by eDNA metabarcoding. Inevitably, there will be a cost-benefit trade-off between using imperfect broad metabarcoding assays for monitoring unknown invaders, and expending resources on the development and application of eDNA tools targeting specific known NNS.

In some cases, resource managers might be interested in tracking invasions using haplotype data [97]. Here, we showed that eDNA metabarcoding with broad-target primers resolves broad scale patterns of haplotype diversity (Fig. 3). However, fine-scale genetic variation was not recovered in our study, indicating that targeted eDNA amplicon sequencing [48] might be more appropriate when this level of genetic data is required. As with biodiversity incidence data, the management objectives for a given NNS determine how haplotype sequencing should be implemented. If large numbers of tissue samples can be easily collected and there are sufficient resources, then sequencing the tissue directly might be more appropriate. In contrast if a broad scale analysis across a larger or difficult-to-sample area is preferred resolving haplotype data from eDNA metabarcoding might be preferable. Overall, eDNA based techniques show great potential for NNS detection but for our target taxa, we demonstrated that current biodiversity surveys and direct tissue sequencing are more reliable for the detection of NNS and genetic composition. It is important to note that there are several key advantages of eDNA-based methods compared to the other tools used in this work. Firstly, eDNA samples can be collected with minimal training and the sequenced DNA provides an unambiguous identification, provided reference data is available [33, 34]. Secondly, eDNA-based methods can be automated and can scale to a much greater survey effort at reduced cost compared to other methods [98]. Third, the limitations described above concerning the sensitivity of eDNA-based incidence data and lack of resolution of eDNA-based haplotype data can be attributed to the use of metabarcoding with broad-target primers. Reanalysing the samples with metabarcoding primers for more specific groups or using species specific qPCR assays [94] would provide increased sensitivity and accuracy.

Overall we demonstrated how our combined methodological approach can effectively detect spatial and temporal trends of range shifts and genetic differentiation, but also to monitor biodiversity changes of both threatened and NSS. The strengths of eDNA or DNA-based biomonitoring demonstrated here for the detection of range shifting species make them a pragmatic choice for natural resources managers. These tools provide managers with greater sensitivity and accuracy when monitoring biodiversity in human impacted environments.

## Acknowledgements

We thank M. Czachur and T. Grevesse for assistance during field surveys and the Elwandle Node of the South African Environmental Observation Network for hosting L.E.H. and for assistance with logistics. We acknowledge the South African Department of Environmental Affairs and the Department of Agriculture, Forestry and Fisheries for granting a research permit (Ref RES2017/100) to L.E.H. L.E.H. was supported by the Natural Environmental Research Council (grant number NE/L002531/1) and research in South Africa was supported by the Newton Fund (grant number ES/N013913/1).

## Authors Contributions

All authors contributed to the initial study design. L.E.H. collected the samples, generated and analysed the data, prepared all figures and wrote the first draft of the paper. S.P.N. and M.R. advised on the sampling design and subsequent laboratory work. All authors substantially contributed to further manuscript drafts and provided final approval for publication.

## Data Availability

Raw and processed data are available online via Zenodo with the following DOI 10.5281/zenodo.5046379. Raw sequence data associated with analyses can be found under European Nucleotide Archive Project Accession PRJEB38452

## Code Availability

Associated R scripts and intermediate files are available online via Zenodo with the following DOI 10.5281/zenodo.5046379.

## References

[1] Pecl, G.T., Araujo, M.B., Bell, J.D., Blanchard, J., Bonebrake, T.C., Chen, I.C., Clark, T.D., Colwell, R.K., Danielsen, F., Evengard, B., et al. 2017 Biodiversity redistribution under climate change: Impacts on ecosystems and human well-being. Science 355:6332. (doi:10.1126/science.aai9214).

[2] Sunday, J.M., Bates, A.E. & Dulvy, N.K. 2012 Thermal tolerance and the global redistribution of animals. Nature Climate Change 2, 686–690. (doi:10.1038/nclimate1539).

[3] Sunday, J.M., Pecl, G.T., Frusher, S., Hobday, A.J., Hill, N., Holbrook, N.J., Edgar, G.J., Stuart-Smith, R., Barrett, N., Wernberg, T., et al. 2015 Species traits and climate velocity explain geographic range shifts in an ocean-warming hotspot. Ecology Letters 18, 944–953. (doi:https://doi.org/10.1111/ele.12474).

[4] Bax, N., Williamson, A., Aguero, M., Gonzalez, E. & Geeves, W. 2003 Marine invasive alien species: a threat to global biodiversity. Marine Policy 27, 313–323. (doi:10.1016/S0308-597x(03)00041-1).

[5] Molnar, J.L., Gamboa, R.L., Revenga, C. & Spalding, M.D. 2008 Assessing the global threat of invasive species to marine biodiversity. Frontiers in Ecology and the Environment 6, 485–492. (doi:10.1890/070064).

[6] Seebens, H., Bacher, S., Blackburn, T.M., Capinha, C., Dawson, W., Dullinger, S., Genovesi, P., Hulme, P.E., van Kleunen, M., Kühn, I., et al. 2021 Projecting the continental accumulation of alien species through to 2050. Global Change Biology 27, 970–982. (doi:https://doi.org/10.1111/gcb.15333).

[7] Dornelas, M., Gotelli, N.J., Shimadzu, H., Moyes, F., Magurran, A.E. & McGill, B.J. 2019 A balance of winners and losers in the Anthropocene. Ecology Letters 22, 847–854. (doi:https://doi.org/10.1111/ele.13242).

[8] Vilà, M., Basnou, C., Pyšek, P., Josefsson, M., Genovesi, P., Gollasch, S., Nentwig, W., Olenin, S., Roques, A., Roy, D., et al. 2010 How well do we understand the impacts of alien species on ecosystem services? A pan-European, cross-taxa assessment. Frontiers in Ecology and the Environment 8, 135–144. (doi:https://doi.org/10.1890/080083).

[9] Gamfeldt, L., Lefcheck, J.S., Byrnes, J.E.K., Cardinale, B.J., Duffy, J.E. & Griffin, J.N. 2015 Marine biodiversity and ecosystem functioning: what’s known and what’s next? Oikos 124, 252–265. (doi:https://doi.org/10.1111/oik.01549).

[10] Cardinale, B.J., Duffy, J.E., Gonzalez, A., Hooper, D.U., Perrings, C., Venail, P., Narwani, A., Mace, G.M., Tilman, D., Wardle, D.A., et al. 2012 Biodiversity loss and its impact on humanity. Nature 486, 59–67. (doi:10.1038/nature11148).

[11] Mazor, T., Doropoulos, C., Schwarzmueller, F., Gladish, D.W., Kumaran, N., Merkel, K., Di Marco, M. & Gagic, V. 2018 Global mismatch of policy and research on drivers of biodiversity loss. Nature Ecology & Evolution 2, 1071–1074. (doi:10.1038/s41559-018-0563-x).

[12] Diagne, C., Leroy, B., Vaissière, A.-C., Gozlan, R.E., Roiz, D., Jarić, I., Salles, J.-M., Bradshaw, C.J.A. & Courchamp, F. 2021 High and rising economic costs of biological invasions worldwide. Nature In Press. (doi:10.1038/s41586-021-03405-6).

[13] Schindler, S., Staska, B., Adam, M., Rabitsch, W. & Essl, F. 2015 Alien species and public health impacts in Europe: a literature review. NeoBiota 27, 1.

[14] Mazza, G., Tricarico, E., Genovesi, P. & Gherardi, F. 2014 Biological invaders are threats to human health: an overview. Ethology Ecology & Evolution 26, 112–129. (doi:10.1080/03949370.2013.863225).

[15] Williams, S.L., Davidson, I.C., Pasari, J.R., Ashton, G.V., Carlton, J.T., Crafton, R.E., Fontana, R.E., Grosholz, E.D., Miller, A.W., Ruiz, G.M., et al. 2013 Managing Multiple Vectors for Marine Invasions in an Increasingly Connected World. BioScience 63, 952–966. (doi:10.1525/bio.2013.63.12.8).

[16] Katsanevakis, S., Zenetos, A., Belchior, C. & Cardoso, A.C. 2013 Invading European Seas: Assessing pathways of introduction of marine aliens. Ocean & Coastal Management 76, 64–74. (doi:https://doi.org/10.1016/j.ocecoaman.2013.02.024).

[17] Airoldi, L., Turon, X., Perkol-Finkel, S. & Rius, M. 2015 Corridors for aliens but not for natives: effects of marine urban sprawl at a regional scale. Diversity and Distributions 21, 755–768. (doi:10.1111/ddi.12301).

[18] Glasby, T.M., Connell, S.D., Holloway, M.G. & Hewitt, C.L. 2007 Nonindigenous biota on artificial structures: could habitat creation facilitate biological invasions? Marine Biology 151, 887–895. (doi:10.1007/s00227-006-0552-5).

[19] Seebens, H., Blackburn, T.M., Dyer, E.E., Genovesi, P., Hulme, P.E., Jeschke, J.M., Pagad, S., Pyšek, P., Winter, M., Arianoutsou, M., et al. 2017 No saturation in the accumulation of alien species worldwide. Nature Communications 8, 14435. (doi:10.1038/ncomms14435).

[20] Booy, O., Robertson, P.A., Moore, N., Ward, J., Roy, H.E., Adriaens, T., Shaw, R., Van Valkenburg, J., Wyn, G., Bertolino, S., et al. 2020 Using structured eradication feasibility assessment to prioritize the management of new and emerging invasive alien species in Europe. Global Change Biology 26, 6235–6250. (doi:https://doi.org/10.1111/gcb.15280).

[21] Roy, H.E., Bacher, S., Essl, F., Adriaens, T., Aldridge, D.C., Bishop, J.D.D., Blackburn, T.M., Branquart, E., Brodie, J., Carboneras, C., et al. 2019 Developing a list of invasive alien species likely to threaten biodiversity and ecosystems in the European Union. Global Change Biology 25, 1032–1048. (doi:https://doi.org/10.1111/gcb.14527).

[22] Thresher, R.E. & Kuris, A.M. 2004 Options for Managing Invasive Marine Species. Biological Invasions 6, 295–300. (doi:10.1023/B:BINV.0000034598.28718.2e).

[23] Clout, M.N. & Williams, P.A. 2009 Invasive species management: a handbook of principles and techniques, Oxford University Press.

[24] Shackleton, R.T., Adriaens, T., Brundu, G., Dehnen-Schmutz, K., Estévez, R.A., Fried, J., Larson, B.M.H., Liu, S., Marchante, E., Marchante, H., et al. 2019 Stakeholder engagement in the study and management of invasive alien species. Journal of Environmental Management 229, 88–101. (doi:https://doi.org/10.1016/j.jenvman.2018.04.044).

[25] Liordos, V., Kontsiotis, V.J., Georgari, M., Baltzi, K. & Baltzi, I. 2017 Public acceptance of management methods under different human–wildlife conflict scenarios. Science of the Total Environment 579, 685–693. (doi:https://doi.org/10.1016/j.scitotenv.2016.11.040).

[26] Simberloff, D. 2020 Maintenance management and eradication of established aquatic invaders. Hydrobiologia. (doi:10.1007/s10750-020-04352-5).

[27] Pluess, T., Cannon, R., Jarošík, V., Pergl, J., Pyšek, P. & Bacher, S. 2012 When are eradication campaigns successful? A test of common assumptions. Biological Invasions 14, 1365–1378. (doi:10.1007/s10530-011-0160-2).

[28] Giakoumi, S., Katsanevakis, S., Albano, P.G., Azzurro, E., Cardoso, A.C., Cebrian, E., Deidun, A., Edelist, D., Francour, P., Jimenez, C., et al. 2019 Management priorities for marine invasive species. Science of the Total Environment 688, 976–982. (doi:https://doi.org/10.1016/j.scitotenv.2019.06.282).

[29] Darling, J.A., Galil, B.S., Carvalho, G.R., Rius, M., Viard, F. & Piraino, S. 2017 Recommendations for developing and applying genetic tools to assess and manage biological invasions in marine ecosystems. Marine Policy 85, 54–64. (doi:https://doi.org/10.1016/j.marpol.2017.08.014).

[30] Hudson, J., Castilla, J.C., Teske, P.R., Beheregaray, L.B., Haigh, I.D., McQuaid, C.D. & Rius, M. 2021 Genomics-informed models reveal extensive stretches of coastline under threat by an ecologically dominant invasive species. Proceedings of the National Academy of Sciences 118, e2022169118. (doi:10.1073/pnas.2022169118).

[31] Leung, B., Lodge, D.M., Finnoff, D., Shogren, J.F., Lewis, M.A. & Lamberti, G. 2002 An ounce of prevention or a pound of cure: bioeconomic risk analysis of invasive species. Proceedings of the Royal Society of London. Series B: Biological Sciences 269, 2407–2413. (doi:doi:10.1098/rspb.2002.2179).

[32] Holman, L.E., de Bruyn, M., Creer, S., Carvalho, G., Robidart, J. & Rius, M. 2019 Detection of introduced and resident marine species using environmental DNA metabarcoding of sediment and water. Scientific Reports 9, 11559. (doi:10.1038/s41598-019-47899-7).

[33] Grey, E.K., Bernatchez, L., Cassey, P., Deiner, K., Deveney, M., Howland, K.L., Lacoursiere-Roussel, A., Leong, S.C.Y., Li, Y., Olds, B., et al. 2018 Effects of sampling effort on biodiversity patterns estimated from environmental DNA metabarcoding surveys. Scientific Reports 8, 8843. (doi:10.1038/s41598-018-27048-2).

[34] Rey, A., Basurko, O.C. & Rodriguez-Ezpeleta, N. 2020 Considerations for metabarcoding-based port biological baseline surveys aimed at marine nonindigenous species monitoring and risk assessments. Ecology and Evolution 10, 2452–2465. (doi:https://doi.org/10.1002/ece3.6071).

[35] Duarte, S., Vieira, P.E., Lavrador, A.S. & Costa, F.O. 2021 Status and prospects of marine NIS detection and monitoring through (e)DNA metabarcoding. Science of the Total Environment 751, 141729. (doi:https://doi.org/10.1016/j.scitotenv.2020.141729).

[36] Deiner, K., Bik, H.M., Machler, E., Seymour, M., Lacoursiere-Roussel, A., Altermatt, F., Creer, S., Bista, I., Lodge, D.M., de Vere, N., et al. 2017 Environmental DNA metabarcoding: Transforming how we survey animal and plant communities. Molecular Ecology 26, 5872–5895. (doi:10.1111/mec.14350).

[37] Fediajevaite, J., Priestley, V., Arnold, R. & Savolainen, V. 2021 Meta-analysis shows that environmental DNA outperforms traditional surveys, but warrants better reporting standards. Ecology and Evolution In Press. (doi:https://doi.org/10.1002/ece3.7382).

[38] Hudson, J., Johannesson, K., McQuaid, C.D. & Rius, M. 2020 Secondary contacts and genetic admixture shape colonization by an amphiatlantic epibenthic invertebrate. Evolutionary Applications 13, 600–612. (doi:https://doi.org/10.1111/eva.12893).

[39] Brown, J.E. & Stepien, C.A. 2009 Invasion genetics of the Eurasian round goby in North America: tracing sources and spread patterns. Molecular Ecology 18, 64–79. (doi:https://doi.org/10.1111/j.1365-294X.2008.04014.x).

[40] Jeffery, N.W., DiBacco, C., Van Wyngaarden, M., Hamilton, L.C., Stanley, R.R.E., Bernier, R., FitzGerald, J., Matheson, K., McKenzie, C.H., Nadukkalam Ravindran, P., et al. 2017 RAD sequencing reveals genomewide divergence between independent invasions of the European green crab (Carcinus maenas) in the Northwest Atlantic. Ecology and Evolution 7, 2513–2524. (doi:https://doi.org/10.1002/ece3.2872).

[41] Russell, J.C., Miller, S.D., Harper, G.A., MacInnes, H.E., Wylie, M.J. & Fewster, R.M. 2010 Survivors or reinvaders? Using genetic assignment to identify invasive pests following eradication. Biological Invasions 12, 1747–1757. (doi:10.1007/s10530-009-9586-1).

[42] Tsuji, S., Miya, M., Ushio, M., Sato, H., Minamoto, T. & Yamanaka, H. 2020 Evaluating intraspecific genetic diversity using environmental DNA and denoising approach: A case study using tank water. Environmental DNA 2, 42–52. (doi:https://doi.org/10.1002/edn3.44).

[43] Holman, L.E., Hollenbeck, C.M., Ashton, T.J. & Johnston, I.A. 2019 Demonstration of the Use of Environmental DNA for the Non-Invasive Genotyping of a Bivalve Mollusk, the European Flat Oyster (Ostrea edulis). Frontiers in Genetics 10, 1159. (doi:10.3389/fgene.2019.01159).

[44] Sigsgaard, E.E., Jensen, M.R., Winkelmann, I.E., Møller, P.R., Hansen, M.M. & Thomsen, P.F. 2020 Population-level inferences from environmental DNA—Current status and future perspectives. Evolutionary Applications 13, 245–262. (doi:https://doi.org/10.1111/eva.12882).

[45] Turon, X., Antich, A., Palacín, C., Præbel, K. & Wangensteen, O.S. 2020 From metabarcoding to metaphylogeography: separating the wheat from the chaff. Ecological Applications 30, e02036. (doi:https://doi.org/10.1002/eap.2036).

[46] Adams, C.I., Knapp, M., Gemmell, N.J., Jeunen, G.-J., Bunce, M., Lamare, M.D. & Taylor, H.R. 2019 Beyond biodiversity: can environmental DNA (eDNA) cut it as a population genetics tool? Genes 10, 192.

[47] Andres, K.J., Sethi, S.A., Lodge, D.M. & Andrés, J. 2021 Nuclear eDNA estimates population allele frequencies and abundance in experimental mesocosms and field samples. Molecular Ecology n/a. (doi:https://doi.org/10.1111/mec.15765).

[48] Sigsgaard, E.E., Nielsen, I.B., Bach, S.S., Lorenzen, E.D., Robinson, D.P., Knudsen, S.W., Pedersen, M.W., Jaidah, M., Orlando, L., Willerslev, E., et al. 2017 Population characteristics of a large whale shark aggregation inferred from seawater environmental DNA. Nature Ecology & Evolution 1. (doi:UNSP 0004 10.1038/s41559-016-0004).

[49] Weitemier, K., Penaluna, B.E., Hauck, L.L., Longway, L.J., Garcia, T. & Cronn, R. 2021 Estimating the genetic diversity of Pacific salmon and trout using multigene eDNA metabarcoding. Molecular Ecology In Press. (doi:https://doi.org/10.1111/mec.15811).

[50] Uchii, K., Doi, H. & Minamoto, T. 2016 A novel environmental DNA approach to quantify the cryptic invasion of non-native genotypes. Molecular Ecology Resourses 16, 415–422. (doi:10.1111/1755-0998.12460).

[51] Griffiths, C.L., Robinson, T.B., Lange, L. & Mead, A. 2010 Marine biodiversity in South Africa: an evaluation of current states of knowledge. PLoS One 5, e12008.

[52] Rius, M., Clusella-Trullas, S., McQuaid, C.D., Navarro, R.A., Griffiths, C.L., Matthee, C.A., von der Heyden, S. & Turon, X. 2014 Range expansions across ecoregions: interactions of climate change, physiology and genetic diversity. Global Ecology and Biogeography 23, 76–88. (doi:10.1111/geb.12105).

[53] Robinson, T.B., Griffiths, C.L., McQuaid, C.D. & Rius, M. 2005 Marine alien species of South Africa — status and impacts. African Journal of Marine Science 27, 297–306. (doi:10.2989/18142320509504088).

[54] Arenas, F., Bishop, J.D.D., Carlton, J.T., Dyrynda, P.J., Farnham, W.F., Gonzalez, D.J., Jacobs, M.W., Lambert, C., Lambert, G., Nielsen, S.E., et al. 2006 Alien species and other notable records from a rapid assessment survey of marinas on the south coast of England. Journal of the Marine Biological Association of the United Kingdom 86, 1329–1337. (doi:10.1017/S0025315406014354).

[55] Millar, R.H. 1962 Further descriptions of South African ascidians, South African Museum.

[56] Michaelsen, W. & Stephenson, T. 1934 The ascidians of the Cape Province of South Africa. Transactions of the Royal Society of South Africa 22, 129–163.

[57] Monniot, C., Monniot, F., Griffiths, C.L. & Schleyer, M. 2001 South African ascidians. Annals of the South African Museum 108, 1–141.

[58] Zhan, A., Briski, E., Bock, D.G., Ghabooli, S. & MacIsaac, H.J. 2015 Ascidians as models for studying invasion success. Marine Biology 162, 2449–2470. (doi:10.1007/s00227-015-2734-5).

[59] Svane, I. & Young, C.M. 1989 The ecology and behaviour of ascidian larvae. Oceanography and Marine Biology: an Annual Review 27, 45–90.

[60] Millar, R. 1955 A collection of ascidians from South Africa. Proceedings of the Zoological Society of London 125, 169–221.

[61] Millar, R. 1964 South African ascidians collected by Th. Mortensen, with some additional material. Videnskabelige Meddelelser Fra Dansk Naturhistorisk Forening 127, 159–180.

[62] Nydam, M.L. & Harrison, R.G. 2007 Genealogical relationships within and among shallow-water Ciona species (Ascidiacea). Marine Biology 151, 1839–1847. (doi:10.1007/s00227-007-0617-0).

[63] Tarjuelo, I., Posada, D., Crandall, K., Pascual, M. & Turon, X. 2001 Cryptic species of Clavelina (Ascidiacea) in two different habitats: harbours and rocky littoral zones in the northwestern Mediterranean. Marine Biology 139, 455–462. (doi:10.1007/s002270100587).

[64] Steinke, D., Prosser, S.W.J. & Hebert, P.D.N. 2016 DNA Barcoding of Marine Metazoans. In Marine Genomics: Methods and Protocols (ed. S.J. Bourlat), pp. 155–168. New York, NY, Springer New York.

[65] Holman, L.E., de Bruyn, M., Creer, S., Carvalho, G., Robidart, J. & Rius, M. 2021 Animals, protists and bacteria share marine biogeographic patterns. Nature Ecology & Evolution 5, 738–746. (doi:https://doi.org/10.1038/s41559-021-01439-7).

[66] Spens, J., Evans, A.R., Halfmaerten, D., Knudsen, S.W., Sengupta, M.E., Mak, S.S.T., Sigsgaard, E.E., Hellström, M. & Yu, D. 2017 Comparison of capture and storage methods for aqueous macrobial eDNA using an optimized extraction protocol: advantage of enclosed filter. Methods in Ecology and Evolution 8, 635–645. (doi:10.1111/2041-210x.12683).

[67] Leray, M., Yang, J.Y., Meyer, C.P., Mills, S.C., Agudelo, N., Ranwez, V., Boehm, J.T. & Machida, R.J. 2013 A new versatile primer set targeting a short fragment of the mitochondrial COI region for metabarcoding metazoan diversity: application for characterizing coral reef fish gut contents. Frontiers in Zoology 10, 34. (doi:10.1186/1742-9994-10-34).

[68] Zhan, A., Hulák, M., Sylvester, F., Huang, X., Adebayo, A.A., Abbott, C.L., Adamowicz, S.J., Heath, D.D., Cristescu, M.E. & MacIsaac, H.J. 2013 High sensitivity of 454 pyrosequencing for detection of rare species in aquatic communities. Methods in Ecology and Evolution 4, 558–565. (doi:10.1111/2041-210x.12037).

[69] Martin, M. 2011 Cutadapt removes adapter sequences from high-throughput sequencing reads. EMBnet Journal 17, 10–12.

[70] Callahan, B.J., McMurdie, P.J., Rosen, M.J., Han, A.W., Johnson, A.J. & Holmes, S.P. 2016 DADA2: High-resolution sample inference from Illumina amplicon data. Nature Methods 13, 581–583. (doi:10.1038/nmeth.3869).

[71] R_Core_Team. 2021 R: a language and environment for statistical computing. (Vienna, Austria, R Foundation for Statistical Computing.

[72] Antich, A., Palacin, C., Wangensteen, O.S. & Turon, X. 2021 To denoise or to cluster, that is not the question: optimizing pipelines for COI metabarcoding and metaphylogeography. BMC Bioinformatics 22, 177. (doi:10.1186/s12859-021-04115-6).

[73] Edgar, R.C. 2013 UPARSE: highly accurate OTU sequences from microbial amplicon reads. Nature Methods 10, 996–998. (doi:10.1038/nmeth.2604).

[74] Rognes, T., Flouri, T., Nichols, B., Quince, C. & Mahe, F. 2016 VSEARCH: a versatile open source tool for metagenomics. PeerJ 4, e2584. (doi:10.7717/peerj.2584).

[75] Charif, D. & Lobry, J.R. 2007 SeqinR 1.0-2: a contributed package to the R project for statistical computing devoted to biological sequences retrieval and analysis. In Structural approaches to sequence evolution (pp. 207–232, Springer.

[76] Paradis, E. 2010 pegas: an R package for population genetics with an integrated– modular approach. Bioinformatics 26, 419–420.

[77] Bandelt, H.-J., Forster, P. & Röhl, A. 1999 Median-joining networks for inferring intraspecific phylogenies. Molecular Biology and Evolution 16, 37–48.

[78] Leigh, J.W. & Bryant, D. 2015 popart: full-feature software for haplotype network construction. Methods in Ecology and Evolution 6, 1110–1116.

[79] Kamvar, Z.N., Tabima, J.F. & Grünwald, N.J. 2014 Poppr: an R package for genetic analysis of populations with clonal, partially clonal, and/or sexual reproduction. PeerJ 2, e281.

[80] Sorte, C.J.B., Williams, S.L. & Carlton, J.T. 2010 Marine range shifts and species introductions: comparative spread rates and community impacts. Global Ecology and Biogeography 19, 303–316. (doi:https://doi.org/10.1111/j.1466-8238.2009.00519.x).

[81] Grosholz, E.D. 1996 Contrasting Rates of Spread for Introduced Species in Terrestrial and Marine Systems. Ecology 77, 1680–1686. (doi:https://doi.org/10.2307/2265773).

[82] Dumont, C.P., Gaymer, C.F. & Thiel, M. 2011 Predation contributes to invasion resistance of benthic communities against the non-indigenous tunicate Ciona intestinalis. Biological Invasions 13, 2023–2034. (doi:10.1007/s10530-011-0018-7).

[83] Rius, M., Potter, E.E., Aguirre, J.D. & Stachowicz, J.J. 2014 Mechanisms of biotic resistance across complex life cycles. Journal of Animal Ecology 83, 296–305.

[84] Pineda, M.C., Turon, X., Pérez-Portela, R. & López-Legentil, S. 2016 Stable populations in unstable habitats: temporal genetic structure of the introduced ascidian Styela plicata in North Carolina. Marine Biology 163, 59. (doi:10.1007/s00227-016-2829-7).

[85] Pérez-Portela, R., Turon, X. & Bishop, J. 2012 Bottlenecks and loss of genetic diversity: spatio-temporal patterns of genetic structure in an ascidian recently introduced in Europe. Marine Ecology Progress Series 451, 93–105.

[86] Pineda, M.-C., Lorente, B., López-Legentil, S., Palacin, C. & Turon, X. 2016 Stochasticity in space, persistence in time: genetic heterogeneity in harbour populations of the introduced ascidian Styela plicata. PeerJ 4, e2158.

[87] Haye, P.A., Turon, X. & Segovia, N.I. 2021 Time or Space? Relative Importance of Geographic Distribution and Interannual Variation in Three Lineages of the Ascidian Pyura chilensis in the Southeast Pacific Coast. Frontiers in Marine Science 8, 413.

[88] Rius, M., Pineda, M.C. & Turon, X. 2009 Population dynamics and life cycle of the introduced ascidian Microcosmus squamiger in the Mediterranean Sea. Biological Invasions 11, 2181–2194. (doi:10.1007/s10530-008-9375-2).

[89] Pineda, M.C., López-Legentil, S. & Turon, X. 2013 Year-round reproduction in a seasonal sea: biological cycle of the introduced ascidian Styela plicata in the Western Mediterranean. Marine Biology 160, 221–230. (doi:10.1007/s00227-012-2082-7).

[90] De Caralt, S., López-Legentil, S., Tarjuelo, I., Uriz, M.J. & Turon, X. 2002 Contrasting biological traits of Clavelina lepadiformis (Ascidiacea) populations from inside and outside harbours in the western Mediterranean. Marine Ecology Progress Series 244, 125–137.

[91] Caputi, L., Toscano, F., Arienzo, M., Ferrara, L., Procaccini, G. & Sordino, P. 2019 Temporal correlation of population composition and environmental variables in the marine invader Ciona robusta. Marine Ecology 40, e12543. (doi:https://doi.org/10.1111/maec.12543).

[92] Chang, A.L., Brown, C.W., Crooks, J.A. & Ruiz, G.M. 2018 Dry and wet periods drive rapid shifts in community assembly in an estuarine ecosystem. Global Change Biology 24, e627–e642. (doi:https://doi.org/10.1111/gcb.13972).

[93] Blackman, R.C., Ling, K.K.S., Harper, L.R., Shum, P., Hanfling, B. & Lawson-Handley, L. 2020 Targeted and passive environmental DNA approaches outperform established methods for detection of quagga mussels, Dreissena rostriformis bugensis in flowing water. Ecology and Evolution 10, 13248–13259. (doi:10.1002/ece3.6921).

[94] Harper, L.R., Lawson Handley, L., Hahn, C., Boonham, N., Rees, H.C., Gough, K.C., Lewis, E., Adams, I.P., Brotherton, P., Phillips, S., et al. 2018 Needle in a haystack? A comparison of eDNA metabarcoding and targeted qPCR for detection of the great crested newt (Triturus cristatus). Ecology and Evolution 8, 6330–6341. (doi:https://doi.org/10.1002/ece3.4013).

[95] Scott, R., Zhan, A., Brown, E.A., Chain, F.J.J., Cristescu, M.E., Gras, R. & MacIsaac, H.J. 2018 Optimization and performance testing of a sequence processing pipeline applied to detection of nonindigenous species. Evolutionary Applications 11, 891–905. (doi:10.1111/eva.12604).

[96] Holman, L.E., Chng, Y. & Rius, M. 2021 How does eDNA decay affect metabarcoding experiments? Environmental DNA In Press.

[97] Darling, J.A. 2015 Genetic studies of aquatic biological invasions: closing the gap between research and management. Biological Invasions 17, 951–971. (doi:10.1007/s10530-014-0726-x).

[98] Gold, Z., Sprague, J., Kushner, D.J., Zerecero, E. & Barber, P.H. 2021 eDNA metabarcoding as a biomonitoring tool for marine protected areas. PLoS One 16(2), e0238557. (doi:10.1101/2020.08.20.258889).

