## Supplementary Information for "Managing human mediated range shifts: understanding spatial, temporal and genetic variation in marine non-native species"

**Supplementary Table 1**
*Sampling sites, latitude, longitude and date of sampling for non-native ascidian species at South African sites used in biodiversity and genetic surveys.*

| ***Site Name*** | **Site Code** | **Latitude (S)** | **Longitude (E)** | **Sampling Date** |
|---|---|---|---|---|
| *Saldanha Bay* | SB | -33.018908 | 17.950512 | 14-Oct-17 |
| *Club Mykonos Marina* | SM | -33.046315 | 18.04028 | 15-Oct-17 |
| *V&A Waterfront Marina* | TB | -33.909055 | 18.419893 | 18-Oct-17 |
| *Hout Bay Yacht Club* | HB | -34.049767 | 18.348042 | 19-Oct-17 |
| *Mossel Bay Marina* | MB | -34.178386 | 22.144918 | 22-Oct-17 |
| *Knysna* | KN | -34.041086 | 23.043592 | 24-Oct-17 |
| *Port Elizabeth Marina* | PE | -33.96692 | 25.634461 | 27-Oct-17 |
| *Bushman's River (Kenton Marina)* | BR | -33.679558 | 26.655841 | 30-Oct-17 |
| *Port Alfred Marina* | PA | -33.593546 | 26.892084 | 31-Oct-17 |
| *East London Marina* | EL | -33.024163 | 27.896269 | 02-Nov-17 |
| *Durban Marina* | DU | -29.862663 | 31.021903 | 08-Nov-17 |
| *Richards Bay Marina* | RB | -28.793908 | 32.079184 | 10-Nov-17 |

### Supplementary Table 2

***a*** *PCR conditions for each species with time shown in minutes and temperature in degrees Celsius.* ***b*** *Sources and sequences (5' to 3' orientation) of primers used in cytochrome c oxidase subunit I sequencing of South African non-native ascidian species, further publication details for primers are provided in the main manuscript references, digital object identifiers are provided in the table*

**a**

| ***Species*** | ***Denaturation*** | | ***Cycles*** | | | | | | | ***Final Extension*** | |
|---|---|---|---|---|---|---|---|---|---|---|---|
| | **Temp** | **Time** | **Temp** | **Time** | **Temp** | **Time** | **Temp** | **Time** | **Cycles** | **Temp** | **Time** |
| ***Ciona intestinalis*** | 95 | 10:00 | 95 | 0:50 | 48 | 0:30 | 72 | 1:00 | 37 | 72 | 7:00 |
| ***Clavelina lepadiformis*** | 95 | 10:00 | 95 | 0:30 | 55 | 0:30 | 72 | 1:00 | 37 | 72 | 7:00 |
| ***Microcosmus squamiger*** | 95 | 10:00 | 95 | 0:30 | 50 | 0:30 | 72 | 1:00 | 35 | 72 | 7:00 |
| ***Styela plicata*** | 95 | 10:00 | 95 | 0:30 | 50 | 0:30 | 72 | 1:00 | 35 | 72 | 7:00 |

**b**

| **Species** | **PrimerF** | **Forward Sequence 5' – 3'** | **PrimerR** | **Reverse Sequence 5' – 3'** | **Source** | **DOI** |
|---|---|---|---|---|---|---|
| ***Ciona intestinalis*** | NydamHarrisF | GAGTAAGAACTGGRTGRACAGTTTAYCCTCC | NydamHarrisR | ATTAAAACTTAATCTAGTAAAAAGAGGRRATCAATGG | Nydam & Harrison, 2010 | 10.1007/s00227-007-0617-0 |
| ***Clavelina lepadiformis*** | Clav2001F | GTACTGAGCTTTCACAAACTGGGCAAT | Clav2001R | TGAAAAAGAATAGGATCTCTCCTTCC | Tarjuelo et al. 2001 | 10.1007/s002270100587 |
| ***Microcosmus squamiger*** | Mc_F | CCTCTGGGTGCCTAAAAAATCA | Mc_R | AGATGTGAATTAAGCGTTGGC | This work | - |
| ***Styela plicata*** | TunF1 | TDTCAACDAATCATAARGATATTRG | TunR1 | TAAACYTCAGGATGTCYAAARAAYCA | Steinke et al. | 10.1007/978-1-4939-3774-5_10 |

**Supplementary Table 3**

*Analysis of molecular variance (AMOVA) partitioning variance found in COI sequences by sampling occasion (year), site, and within samples. Model outputs are shown for all species.*

| *Ciona robusta* | Df | SumSq | Mean Sq | P value | % variance |
|---|---|---|---|---|---|
| **Between sampling years** | 1 | 29.40128 | 29.401285 | 0.20879121 | 5.879724 |
| **Between sites within year** | 9 | 135.00643 | 15.000715 | 0.000999 | 27.8254 |
| **Within samples** | 264 | 348.71228 | 1.32088 | 0.000999 | 66.294876 |
| **Total** | 274 | 513.12 | 1.872701 | | 100 |

| *Clavelina lepadiformis* | Df | SumSq | Mean Sq | P value | % variance |
|---|---|---|---|---|---|
| **Between sampling years** | 1 | 6.30415 | 6.3041504 | 0.18481519 | 1.489209 |
| **Between sites within year** | 17 | 62.33149 | 3.6665583 | 0.000999 | 19.434993 |
| **Within samples** | 452 | 235.94185 | 0.5219953 | 0.000999 | 79.075799 |
| **Total** | 470 | 304.57749 | 0.6480372 | | 100 |

| *Styela plicata* | Df | SumSq | Mean Sq | P value | % variance |
|---|---|---|---|---|---|
| **Between sampling years** | 1 | 15.92812 | 15.928116 | 0.6953047 | -8.706159 |
| **Between sites within year** | 7 | 674.5894 | 96.369914 | 0.000999 | 42.221948 |
| **Within samples** | 219 | 1244.70179 | 5.68357 | 0.000999 | 66.484211 |
| **Total** | 227 | 1935.2193 | 8.525195 | | 100 |

| *Microcosmus squamiger* | Df | SumSq | Mean Sq | P value | % variance |
|---|---|---|---|---|---|
| **Between sampling years** | 1 | 0.867717 | 0.867717 | 0.98201798 | -1.065651 |
| **Between sites within year** | 10 | 29.099302 | 2.90993 | 0.02697303 | 3.166193 |
| **Within samples** | 234 | 413.081762 | 1.765307 | 0.04295704 | 97.899458 |
| **Total** | 245 | 443.04878 | 1.808362 | | 100 |

**Supplementary Note 1**
*Comparisons between rapid assessment surveys and eDNA metabarcoding derived relative abundance measures for ascidians along the coast of South Africa.*

In order to explore if there was a relationship between the abundance assessments from the rapid assessment surveys and the eDNA metabarcoding surveys, the 18S & COI metabarcoding data was parsed as in the main manuscript for the nine non-native ascidians found along the South African coastline [following [1]]. These species are *Ascidia sydneiensis, Ascidiella aspersa, Asterocarpa humilis, Botryllus schlosseri, Ciona robusta, Clavelina lepadiformis, Diplosoma listerianum, Microcosmus squamiger, Styela plicata*. For the 18S data, sequences could be assigned to *A. aspersa, A. humilis, B. schlosseri, M. squamiger, S. plicata*. For the COI data, sequences could be assigned to all species except *A. sydneiensis* and *A. humilis*. All zero abundance measures were removed from the analysis. Linear models of log10 transformed read proportions against categorical rapid assessment for all species combined were used to assess the relationship between metabarcoding reads and rapid assessment abundance (see Supplementary Note 1 Table 1 below). No significant relationship was found between metabarcoding reads and rapid assessment abundance in either the 18S or COI data ($p > 0.05$).

**Supplementary Note 1 Table 1**
*Table detailing model outputs for ordinary least squares regression models between rapid assessment survey abundance and log10 eDNA metabarcoding relative read abundance. Data is shown for both 18S and COI.*

| **18S** | **Estimate** | **Std. Error** | **t value** | **p value** |
|---|---|---|---|---|
| **(Intercept)** | -3.2924 | 0.399 | -8.252 | 3.51E-08 |
| **Rapid Assessment** | 0.4187 | 0.2038 | 2.055 | 0.052 |
| **F value** | 4.221 | 1 and 22 Df | | |

| **COI** | **Estimate** | **Std. Error** | **t value** | **p value** |
|---|---|---|---|---|
| **(Intercept)** | -4.7869 | 0.5733 | -8.35 | 5.98E-08 |
| **Rapid Assessment** | 0.2425 | 0.2264 | 1.071 | 0.297 |
| **F value** | 1.147 | 1 and 20 Df | | |

**Supplementary Note 1 Figure 1**

*Plots showing rapid assessment survey abundance against log10 eDNA metabarcoding relative read abundance for 18S (left) and COI (right). Rapid assessment abundance is measured in categories of 1(scarce), 2(common) or 3(dominant) following the main text.*

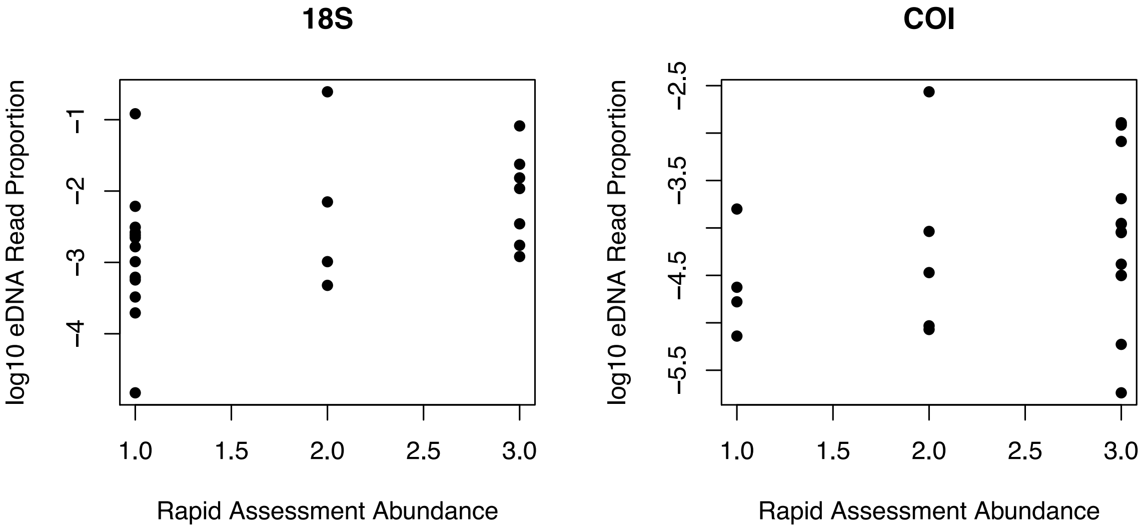

**Supplementary Note 2**
*Analyses of nucleotide and haplotype diversity for non-native ascidians across South Africa.*

In order to explore if there was a significant change in genetic diversity between sampling occasions, data was subset to include only sites where genetic data was collected during both sampling occasions (2009 & 2017) for all species. Nucleotide and haplotypic diversity were calculated using the *nuc.div* and *hap.div* functions from the *pegas* package. Paired samples T-tests were then performed on the diversity estimates to evaluate if there was a significant difference in both haplotype and nucleotide diversity between sampling occasions across the different species: *Ciona robusta; Clavelina lepadiformis; Microcosmus squamiger; Styela plicata.*

**Supplementary Note 2 Figure 1**
*Nucleotide diversity for 2009 and 2017 sampling occasions from sequenced tissue samples. Observations from the same site are connected with dashed red lines, the grey boxes indicate 25% and 7% quartiles with the median value shown by a white dash.* ***a*** *Ciona robusta* ***b*** *Clavelina lepadiformis* ***c*** *Microcosmus squamiger* ***d*** *Styela plicata*

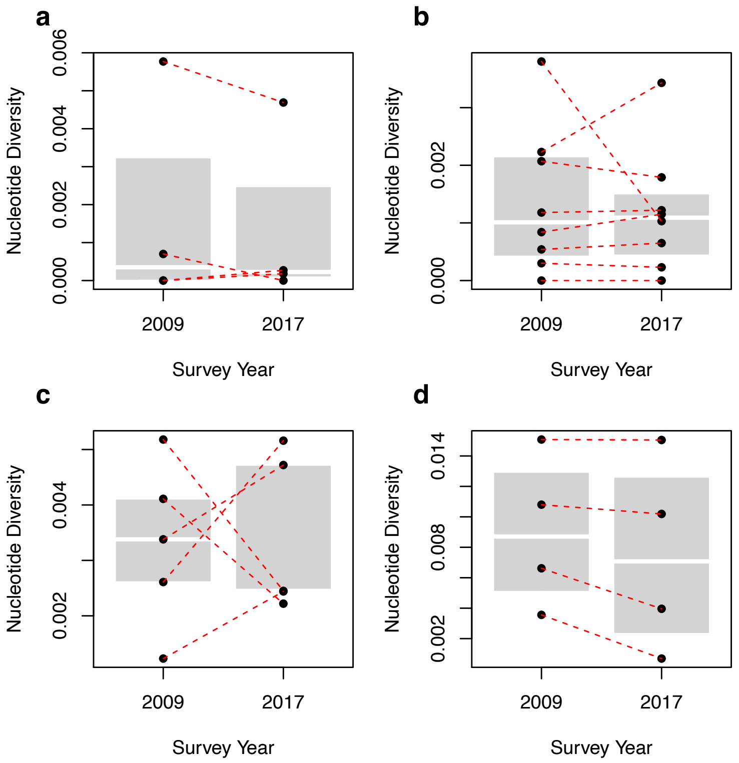

**Supplementary Note 2 Figure 2**
*Haplotype diversity for 2009 and 2017 sampling occasions from sequenced tissue samples. Observations from the same site are connected with dashed red lines, the grey boxes indicate 25% and 7% quartiles with the median value shown by a white dash.* ***a*** *Ciona robusta* ***b*** *Clavelina lepadiformis* ***c*** *Microcosmus squamiger* ***d*** *Styela plicata*

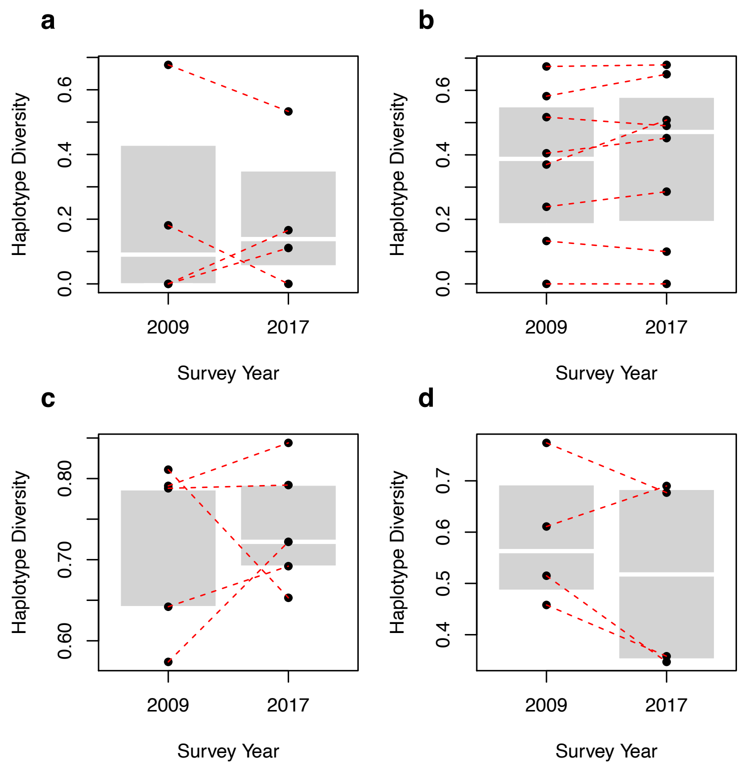


**Supplementary Note 2 Table 1**
*Model output from paired sample T-test conducted on haplotype and nucleotide diversity between sampling occasions. Each row indicates a different model with species and statistic noted for each case. Model t value, degrees of freedom (Df) and p values are shown.*

| Species | Statistic | t value | Df | p value |
|---|---|---|---|---|
| *Ciona robusta* | Haplotype diversity | 0.13646 | 3 | 0.9001 |
| *Ciona robusta* | Nucleotide diversity | 1.0027 | 3 | 0.3899 |
| *Clavelina lepadiformis* | Haplotype diversity | -1.5304 | 7 | 0.1698 |
| *Clavelina lepadiformis* | Nucleotide diversity | 0.45424 | 7 | 0.6634 |
| *Microcosmus squamiger* | Haplotype diversity | -0.3868 | 4 | 0.7186 |
| *Microcosmus squamiger* | Nucleotide diversity | -0.094269 | 4 | 0.9294 |
| *Styela plicata* | Haplotype diversity | 1.3548 | 3 | 0.2685 |
| *Styela plicata* | Nucleotide diversity | 2.1515 | 3 | 0.1205 |

**Supplementary Note 3**

*Evaluation of a raw read mapping approach for the detection of non-native ascidians across South Africa*

Previous work has identified that bioinformatic methods and parameters used in the analysis of DNA metabarcoding data can have an effect on biodiversity metrics [2, 3]. Therefore, a raw read mapping approach was tested as follows. For *Ciona robusta*, *Clavelina lepadiformis, Styela plicata* and *Microcosmus squamiger* all the available cytochrome c oxidase subunit I gene (COI) sequences on NCBI Nucleotide database were downloaded on 11/05/2021. Raw eDNA metabarcoding reads from the COI gene region sequenced across the sites in the main manuscript for each sample were then mapped against the downloaded COI sequences using the --usearch_global setting of vsearch (v2.10.4) with the following parameters *-id 0.97 -strand both -maxhits 1*. The resultant mappings were then summed for each species in each site to produce Table 1 below.

**Supplementary Note 3 Table 1**

*Table showing presence (1) and absence (0) of ascidians in sites across South Africa from eDNA metabarcoding data analysed by mapping to COI regions. Underlined values show detections previously not found when analysing the dataset using a clustering and QC pipeline. Values marked with '!' showed read counts lower than those found in negative control samples. Values marked with '*' correspond with singleton detections that are typically discarded in bioinformatic pipelines. Site abbreviations as in Supplementary Table 1.*

| | SB | SM | TB | HB | MB | KN | PE | BR | PA | EL | DU | RB |
|---|---|---|---|---|---|---|---|---|---|---|---|---|
| *Ciona robusta* | 1 | 0 | 1 | 1* | 0 | 1* | 0 | 0 | 0 | 0 | 0 | 0 |
| *Clavelina lepadiformis* | 1 | 1 | 1 | 1* | 1 | 1 | 1* | 0 | 1 | 0 | 0 | 0 |
| *Microcosmus squamiger* | 0 | 0 | 0 | 0 | 1 | 1 | 1 | 1* | 1 | 1 | 0 | 1 |
| *Styela plicata* | 0 | 1 | 1!* | 1!* | 1 | 1 | 1 | 0 | 0 | 1!* | 1!* | 1 |

The results show several instances of positive detections not found when using a clustering/denoising approach that match up with positive detections from rapid assessment surveys. However, many of these positive detections correspond with one or two reads, which are typically discarded regardless of bioinformatic pipeline. Additionally, one read from a single negative control sample was mapped to *Styela plicata* which suggests extremely low, but non-zero cross-contamination at some stage across the workflow, which is typical of metabarcoding datasets. This shows that while it may be possible to generate more sensitive incidence data by using a raw mapping approach it may not be advisable as the possibility for false positive errors increases.

**Supplementary References**

[1] Rius, M., Clusella-Trullas, S., McQuaid, C.D., Navarro, R.A., Griffiths, C.L., Matthee, C.A., von der Heyden, S. & Turon, X. 2014 Range expansions across ecoregions: interactions of climate change, physiology and genetic diversity. *Global Ecology and Biogeography* **23**, 76-88. (doi:10.1111/geb.12105).
[2] Scott, R., Zhan, A., Brown, E.A., Chain, F.J.J., Cristescu, M.E., Gras, R. & MacIsaac, H.J. 2018 Optimization and performance testing of a sequence processing pipeline applied to detection of nonindigenous species. *Evolutionary Applications* **11**, 891-905. (doi:10.1111/eva.12604).
[3] Antich, A., Palacin, C., Wangensteen, O.S. & Turon, X. 2021 To denoise or to cluster, that is not the question: optimizing pipelines for COI metabarcoding and metaphylogeography. *BMC Bioinformatics* **22**, 177. (doi:10.1186/s12859-021-04115-6).
